# Guidelines for accurate genotyping of SARS-CoV-2 using amplicon-based sequencing of clinical samples

**DOI:** 10.1101/2020.12.01.405738

**Authors:** Slawomir Kubik, Ana Claudia Marques, Xiaobin Xing, Janine Silvery, Claire Bertelli, Flavio De Maio, Spyros Pournaras, Tom Burr, Yannis Duffourd, Helena Siemens, Chakib Alloui, Lin Song, Yvan Wenger, Alexandra Saitta, Morgane Macheret, Ewan W. Smith, Philippe Menu, Marion Brayer, Lars M. Steinmetz, Ali Si-Mohammed, Josiane Chuisseu, Richard Stevens, Pantelis Constantoulakis, Michela Sali, Gilbert Greub, Carsten Tiemann, Vicent Pelechano, Adrian Willig, Zhenyu Xu

## Abstract

**Background:** SARS-CoV-2 genotyping has been instrumental to monitor virus evolution and transmission during the pandemic. The reliability of the information extracted from the genotyping efforts depends on a number of aspects, including the quality of the input material, applied technology and potential laboratory-specific biases. These variables must be monitored to ensure genotype reliability. The current lack of guidelines for SARS-CoV-2 genotyping leads to inclusion of error-containing genome sequences in studies of viral spread and evolution.

**Results:** We used clinical samples and synthetic viral genomes to evaluate the impact of experimental factors, including viral load and sequencing depth, on correct sequence determination using an amplicon-based approach. We found that at least 1000 viral genomes are necessary to confidently detect variants in the genome at frequencies of 10% or higher. The broad applicability of our recommendations was validated in >200 clinical samples from six independent laboratories. The genotypes of clinical isolates with viral load above the recommended threshold cluster by sampling location and period. Our analysis also supports the rise in frequency of 20A.EU1 and 20A.EU2, two recently reported European strains whose dissemination was favoured by travelling during the summer 2020.

**Conclusions:** We present much-needed recommendations for reliable determination of SARS-CoV-2 genome sequence and demonstrate their broad applicability in a large cohort of clinical samples.

## Background

Severe acute respiratory syndrome coronavirus 2 (SARS-CoV-2), a member of the Coronaviridae family, causes a respiratory syndrome and is at the origin of the pandemic that started at the end of 2019 (1-3). Rapid worldwide spread of this pathogen, infecting millions and killing hundreds of thousands, has led to an unprecedented global effort to characterize its genome. Genomic epidemiology, where pathogen genotype information is used to track the spread of a disease, is indispensable for public health monitoring and for high-resolution contact tracing (4,5). For example, genotyping efforts allowed to detect an increase in the prevalence of D614G amino-acid variant in the SARS-CoV-2 Spike protein-encoding gene (23403:A->G) (6,7); and provided evidence for SARS-CoV-2 mink-human transmission (8). Although the clinical relevance of these and similar observations remains controversial (9), virus mutation and cross-species transmission have the potential to change the disease severity and facilitate evasion of the immune response (10) and must be monitored. The unprecedented extent and speed of SARS-CoV-2 genome sequencing since the beginning of the pandemic (*e.g.* 175’000 SARS-CoV-2 genomes deposited as of November 2020 (https://www.gisaid.org) is a testimony of how crucial genotyping has been in attempting to understand and control virus transmission.

Such a large increase in genomic information is inevitably associated with genome submissions of variable quality, depending on laboratory-specific protocol implementation and practices, sample quality, genotyping method and data processing approaches. The genotyping errors introduced as a consequence of these different factors can impact the conclusions of downstream analyses (11-13). Even stringently selected complete genome assemblies obtained from public repositories seem to contain a significant fraction of inaccuracies (14).

Broadly applicable guidelines that take into account the performance metrics of different genotyping approaches are essential to limit the number of inaccurate variant calls in viral sequences submitted to public repositories and used in genomic epidemiology studies. The sensitivity, specificity and limit of detection are dependent of the chosen method (15). Amplicon-based methods are the most widely used approach for SARS-CoV-2 genotyping due to their relative low cost and simplicity (16-22). Despite their widespread use, amplicon-based technologies are associated with source-specific factors that, if not accounted for, can affect the reliability of assigned genotypes (11-14) and impact, for example, the measurements of viral intra-host variability (23). Previous efforts to genotype the Zika virus using amplicon-based approaches suggested that the fraction at which a variant can be confidently separated from technical noise is no lower than 3% even when sufficient input material, sequencing depth and replicates are used (24). However, some of the variants considered in the analysis of SARS-CoV-2 intra-host genome variability were reported at lower fractions (25-27). Furthermore, data generated by different laboratories might contain specific biases, compromising their direct comparison and the validity of downstream analysis (11,12). It is thus essential to provide guidelines and technical recommendations to enable accurate SARS-CoV-2 genotyping and robust comparisons across multiple institutions.

In this work, we aimed to establish guidelines for the implementation of amplicon-based SARS-CoV-2 genotyping. We evaluated the impact of viral load, sequencing depth and coverage uniformity on assay performance using synthetic reference SARS-CoV-2 genomes. To ensure the wide applicability of our conclusions, we performed a multicentre validation study where we benchmarked the same experimental protocol applied to over 200 clinical samples in six independent laboratories across Europe (**Figure 1A**). We demonstrate that SARS-CoV-2 variants with frequencies of 10% or higher can be reproducibly detected with sufficient input material and sequencing depth. Our study provides general recommendations for reliable determination of viral genome sequences using amplicon-based methods for SARS-CoV-2 genotyping.

**Figure 1.**
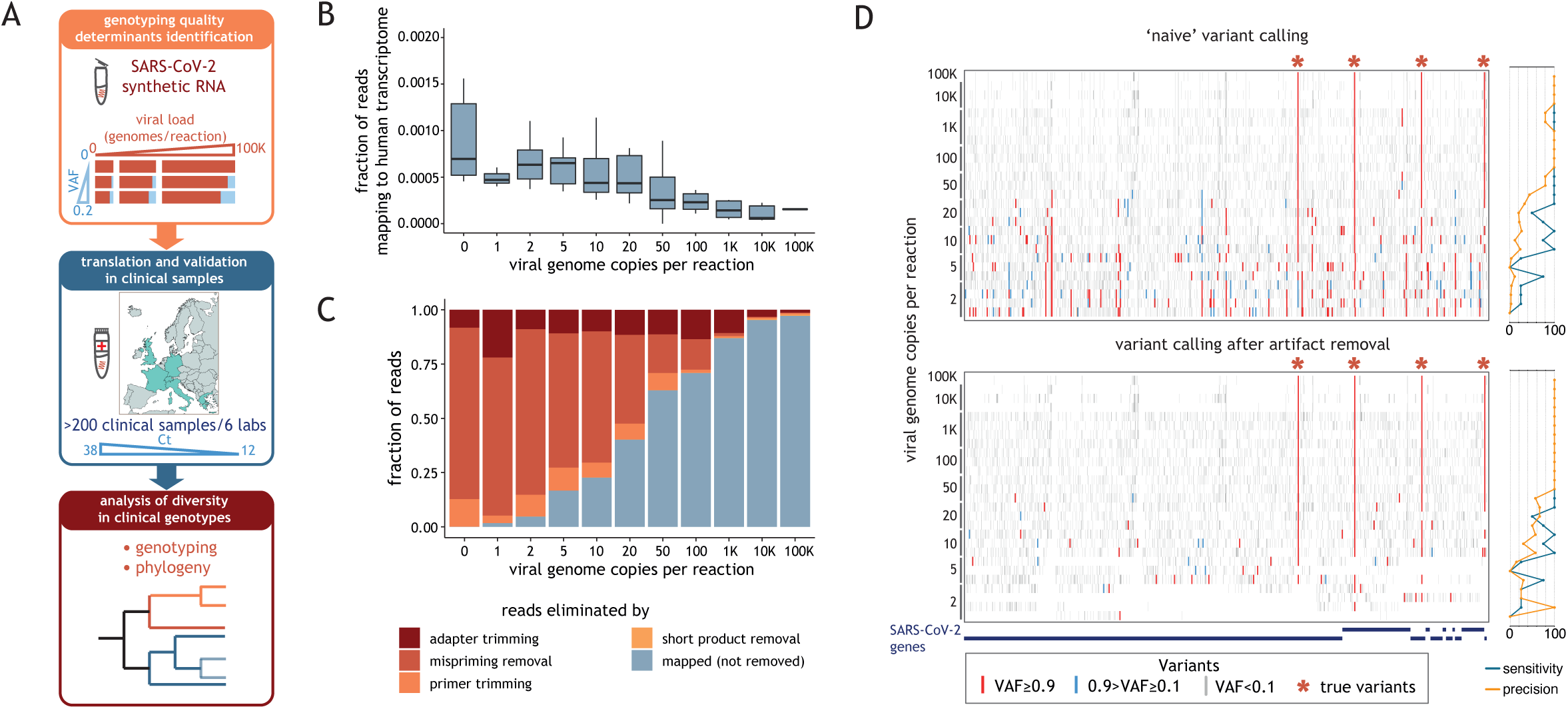
Artifact removal as a prerequisite for reliable variant calling. (A) Schematic representation of the study; In experiments using synthetic SARS-CoV-2 RNA we varied a number of experimental parameters including viral load, VAF and sequencing depth, and determined which of these factors critically impact genotyping quality (top box). We validated these metrics using data obtained from clinical samples, whose viral load is reflected by the Ct value (middle box). We determined the phylogeny of all clinical samples that met our guidelines (bottom box). (B) Distribution of the fraction of raw reads aligning to human transcriptome (y-axis), obtained with STAR aligner, as a function of the number of synthetic viral genome in the sample (x-axis). The horizontal line in the boxplot indicates the median and the whiskers the 5% and 95% quantile (C) Average fraction (from at least 3 replicates) of sequencing reads that mapped to the SARS-CoV-2 genome or were the result of different technical artifacts (y-axis) for samples with varying amounts of synthetic viral genomes (x-axis). (D) Ideogram depicting the location of the variants detected in samples with varying synthetic viral RNA copies (denoted on the left of the ideogram) before (top panel) and after(bottom panel) removal of reads labelled as technical artifacts. Variants with allele fraction below 0.01, between 0.01 and 0.9 and above 0.9 are shown in grey, blue and red, respectively. SARS-CoV-2 variants previously reported (known variants) are marked with asterisks. Plots on the right show sensitivity and precision of the variant calls.

## Results

### Benchmarking of SARS-CoV-2 amplicon-based genotyping using synthetic genome control

The commonly used amplicon-based genotyping method used here is based on generation of 343 partially overlapping amplicons split in two pools, covering positions 36-29844 (99.89%) of the SARS-CoV-2 genome (18). Briefly, viral genomic RNA is converted into cDNA and subsequently amplified by PCR with two sets of primer pairs resulting in two amplicon pools. After combining these pools, amplicons undergo a second PCR step where sequencing adapters are added (**Figure S1A, Additional File 1**).

We first used synthetic RNA controls of the SARS-CoV-2 genome to investigate how the quality of viral genome sequencing data is impacted by the number of viral genome copies in a sample. We spiked 50 ng of reference human RNA with varying amounts of synthetic SARS-CoV-2 genome (1-100’000 genome copies per reaction (g.c.p.r.), the equivalent of ~0.1-10’000 viral genome copies per ml). The synthetic SARS-CoV-2 genome consists of six individual RNA molecules, each ~5 kb in length. We processed and sequenced these samples, in replicates, at a median depth of 1.1M reads. The virtual absence of sequencing reads mapping to the human transcriptome (**Figure 1B**) or genome (**Figure S1B, Additional File 1**) supports the specificity of the assay and alleviates the legal, ethical and technical concerns resulting from the presence of patient sequence information seen in metagenomic or capture-based methods (3,28,29).

We systematically detected and removed sequencing artifacts generated by primer dimers, short uninformative reads, mispriming events and adapter dimers (**Figure S1A, Additional File 1**). The efficiency of the expected product amplification increased with the viral load (**Figure S1C and S1D, Additional File 1**). As expected, the total fraction of effective reads (i.e. reads mapped to SARS-CoV-2 genome after read filtering) correlated with the number of viral copies in the sample (**Figure 1C**). For example, fewer than half of the reads mapped to the reference SARS-CoV-2 genome in samples with 20 or less viral g.c.p.r. (**Figure 1C**). Inclusion of sequencing artifacts resulted in both false-negative and false-positive calls (**Figure S1E, Additional File 1**). This lead to inaccurate variant calling and genotype assignment, as demonstrated in experiments with SARS-CoV-2-Control 1 (hereafter SARS-CoV-2-C1, equivalent to sequence MT007544.1) - a viral genome control carrying 3 Single Nucleotide Variants (SNVs) and a 10-nucleotide deletion distinguishing it from SARS-CoV-2 reference genome (MN 908947.3, SARS-CoV-2 Control 2, hereafter SARS-CoV-2-C2) (**Figure 1D**).

### Guidelines for reliable detection of clonal variants

Coverage breadth (% of genome covered) and depth (number of reads covering each position) are critical determinants of genotyping reliability. We observed that both depended on the number of viral genome copies in the input material. For example, samples containing fewer than 50 SARS-CoV-2 g.c.p.r. displayed only partial genome coverage (**Figure 2A and S2A, Additional File 1**). We aimed to establish the number of sequencing reads required to achieve >10x genome coverage across at least 98% of positions. To this end, we randomly downsampled the number of mapping reads to defined values between 5K and 500K in samples with 10’000 g.c.p.r. These samples were chosen due to low fraction of artifacts (<11% discarded reads) and large number of available replicates (n=15). We determined coverage breadth (**Figure 2B**) and depth (**Figure 2C**), as a function of the number of mapped reads. We found that at least 200K mapped 150 bp paired-end reads are required to ensure 98% genome coverage with an average depth of 683x and with 93% of coverage uniformity (defined as fraction of genome positions falling within a range between 20% and 500% of the median coverage) (**Figure S2B, Additional File 1**). An increase in the number of mapped reads did not translate into any significant improvement in coverage breadth, depth or uniformity (**Figures 2B, 2C** and **S2B, Additional File 1**).

**Figure 2.**
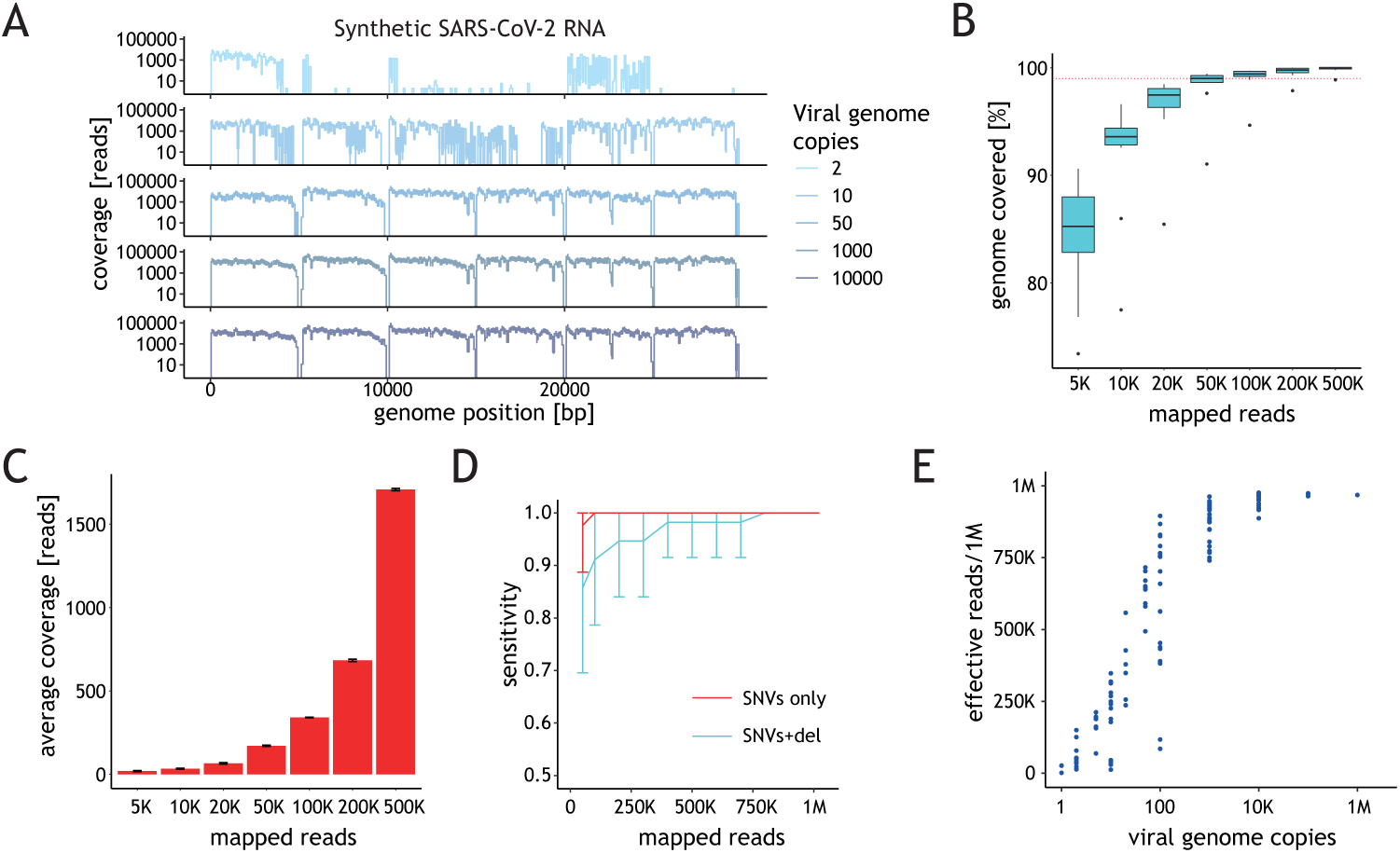
Performance of the assay depends on the amount of starting material. (A) Ideograms depicting the genome coverage (y-axis) for representative samples with varying amount of synthetic viral genomes (x-axis). (B) Distribution of the percentage of the genome covered (y-axis) as a function of the number of mapped reads. Horizontal red dashed line depicts 98% genome coverage. The horizontal line in the boxplot indicates the median and the whiskers the 5% and 95% quantile. (C) Average coverage depth across all synthetic SARS-CoV-2 genome (y-axis) as a function of the number of mapped reads (x-axis) (D) Average sensitivity of variant calling for SNVs (red) or SNV+10-bp indel (cyan) in SARS-CoV-2-C1 (y-axis) as a function of the number of mapped reads based on the results obtained for samples with at least 98% genome coverage breadth. Error bars represent standard deviation. (E) Effective reads per million reads (Y-axis) shown as a function of the viral genome amount (g.c.p.r) in the sample. Each point represents the data for one sample.

Library preparation and sequencing introduce errors that need to be distinguished from true variants to ensure reliable genotyping results. To investigate the impact of sequencing depth on clonal variant calling we used samples containing SARS-CoV-2-C1. We determined that the sensitivity of clonal variant calling achieved 100% at all depths above 50K mapped reads for SNVs (**Figure 2D**). The 10-nucleotide deletion present in SARS-CoV-2-C1 was initially omitted in the analysis as the overlap between the annealing site of one of the amplicon primers and this region of the viral genome strongly decreased the PCR efficiency (**Figure S2C, Additional File 1**). Achieving 100% sensitivity when taking the deletion into account required at least 800K reads (**Figure 2D**). We conclude, that for variants that do not significantly impair PCR amplification, the recommended number of 200K mapped reads is sufficient to reliably call SNVs.

The sequencing depth required to achieve adequate number of mapped reads was inversely proportional to the number of viral genome copies in the sample (**Figure 2E**). For samples with fewer than 1000 viral g.c.p.r. the coverage uniformity (average 51.2%) (**Figure S2D, Additional File 1**) and the percentage of genome covered (average 67.9%) (**Figure S2A, Additional File 1**) were highly variable. This reflects the technical challenges of handling samples with relatively low input material, which can hinder reliable viral genotyping. To ensure confident variant calling we recommend restricting viral genotyping to samples with 1000 or more viral copies g.c.p.r. For these samples the fraction of effective reads in our dataset is relatively stable: at least 740K per million reads (K/M) (910 +/-65K/M on average). For samples with similar or higher viral load, we advise sequencing to a depth of at least 270K reads to achieve the recommended ~200K mapped reads.

### Limits of performance for intra-host variability measurement

Next, we aimed to determine the lowest allele fraction which can be confidently measured using the recommended minimal viral load of 1000 g.c.p.r. We mixed defined amounts of SARS-CoV-2-C1, SARS-CoV-2-C2 and SARS-CoV-2 Control 4 (MT106054.1, hereafter referred to as SARS-CoV-2-C4) to obtain alleles fractions between 0.01 and 0.2 (**Figure 3A**). SARS-CoV-2-C4 has seven SNVs relative to the reference genome, which together with SARS-CoV-2-C1 yielded 10 SNVs per sample in total. We spiked 1000 viral genome copies of these mixes into 50 ng of human RNA, processed and sequenced the samples in 4 replicates for each variant allele frequency (VAF) (median sequencing depth 1.4M reads). To assess the reliability of low frequency variant detection at the recommended 200K, we randomly down-sampled mapped reads to this number and performed variant calling, treating the known SNVs as true positive calls and all other SNVs passing the variant caller filters as false positives generated by the technical noise (**Figure 3B**). Detected variants that we attributed to technical noise had similar average VAF across samples (average VAF=0.018). At this sequencing depth, and for 1000 g.c.p.r., the VAF of >95% of these false-positive variants was lower than the VAF of true-positives in samples with expected variant fractions of 0.1 or higher. The ROC (receiver operating characteristic) curves plotted for samples with different expected variant fractions (**Figure 3C**) showed that at 0.1 expected VAF the separation between the true and false positive calls was within an acceptable range for this type of analysis, with 95% sensitivity and 95% specificity. As expected, the accuracy increased at higher target VAF, with variants present at 0.2 expected VAF being detected with 95% sensitivity and 98% specificity, for example.

**Figure 3.**
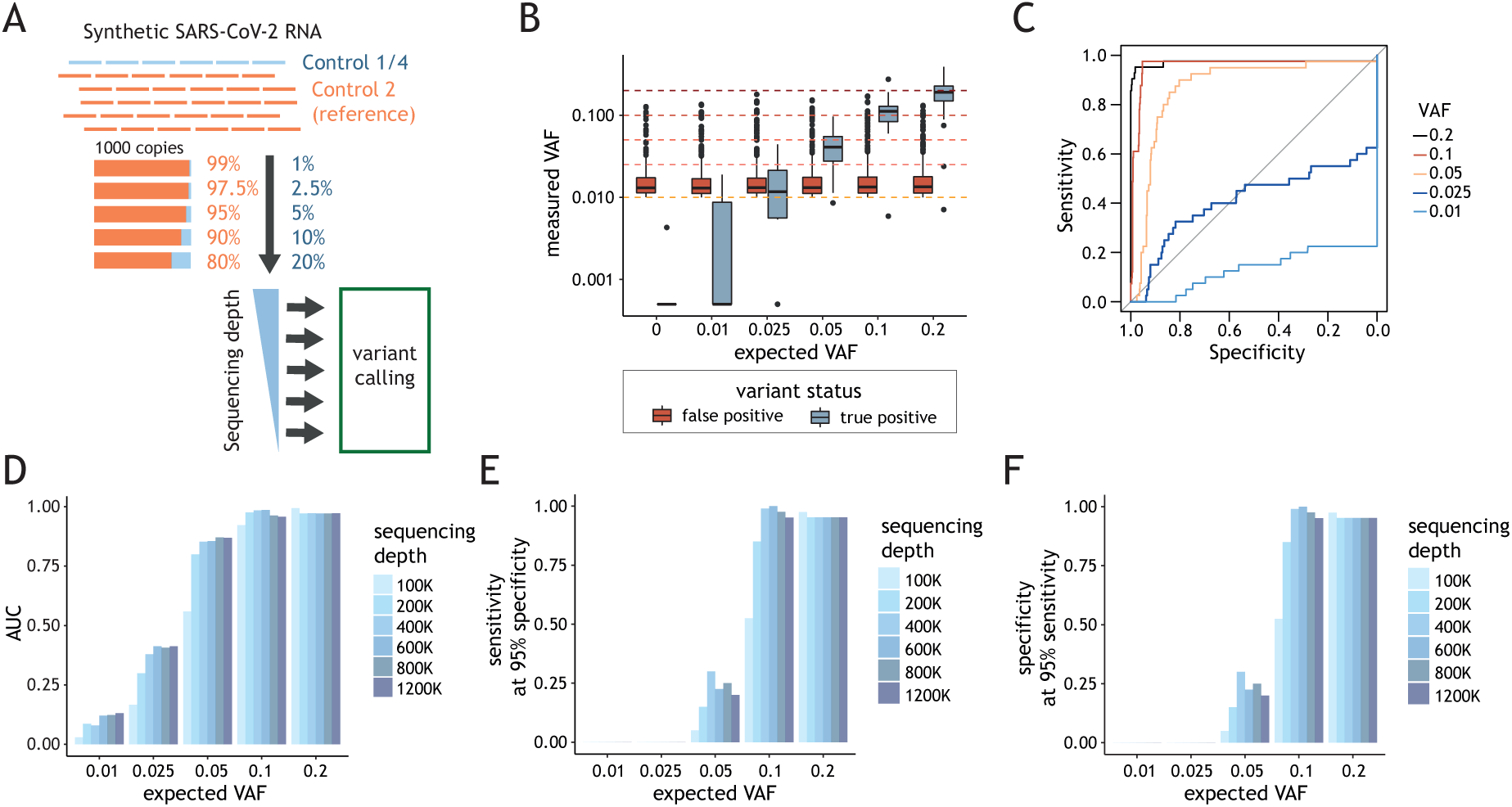
Determination of assay parameters for reliable intra-host variability detection. (A) Schematic representation of the experimental design for the analysis of the intra-host variant calling fraction threshold. Varying amounts of SARS-CoV-2 Control 1 or 4 (blue) were mixed with SARS-CoV-2 synthetic genome reference (Control 2) to obtain desired VAFs. 1000 viral genome copy mixes (g.c.p.r.) were spiked into human RNA. Variant calling was performed for samples sequenced at varying depths. (B) Distribution of variant fraction measured for known (true positives, red) and background (false positives, blue) variants (y-axis) as a function of the expected VAFs in the samples (x-axis). The horizontal line in the boxplot indicates the median and the whiskers the 5% and 95% quantile. (C) Sensitivity (y-axis) as a function of the specificity (x-axis) with VAF value used as a predictor for true variant calls. The ROC curves are colour coded depending on expected VAF of the known variants in each experiment. (D) Area under the ROC curve (AUC) (y-axis) as a function of the expected VAF of the variant (x-axis) at sequencing depth between 100K and 1200K reads. Colour-code for analysis done with samples at different sequencing depth is depicted on the right. (E) Sensitivity at 95% specificity (y-axis) and (F) specificity at 95% sensitivity (y-axis) as a function the expected VAF for the variant (x-axis). Colour-code for analysis done with samples at different sequencing depth is depicted on the right.

To assess the impact of the number of mapped reads on variant calling performance, we repeated variant calling with down-sampled datasets (100K-1.2M reads). As expected, higher numbers of mapped reads resulted in increased sensitivity (**Figure 3D** and **3E**) and specificity (**Figure 3F**) but the increase was modest above 200K reads. Variants present at VAF=0.05 or lower could not be accurately detected in the tested conditions as reflected by low sensitivity and specificity (**Figure 3E** and **3F**).

We performed variant calling on sequencing results obtained with samples bearing 100, 1000 or 10K g.c.p.r. to investigate the impact of viral load on genotyping accuracy (**Figure S3A, Additional File 1**). We observed that using 100 g.c.p.r. resulted in a clear decrease in variant calling reliability, relative to 1000 g.c.p.r. (**Figure S3B and S3C, Additional File 1**). Even if the measured VAF was less variable for samples with higher viral load (**Figure S3D, Additional File 1**), using 10K viral genomes did not further improve the accuracy of variant detection.

In summary, we demonstrated that in order to confidently detect clonal variants, at least 1000 viral genomes are required in the input material and at least 200K mapped reads are recommended. In these conditions, variants present at a 10% or higher frequency can be accurately detected.

### Multicentre study design for assessment of robust SARS-CoV-2 genotyping

Our analysis using synthetic genome controls provided comprehensive information on how viral genome counts impacts several sequencing quality metrics and allowed us to define guidelines to ensure sensitive and specific SARS-CoV-2 amplicon-based genotyping for samples with known viral load. However, the exact number of viral genomes is unknown for most clinical isolates. To address this limitation, we sought to translate our guidelines and make them broadly applicable to the analysis of clinical samples.

Towards this end, we designed a multicentre study involving six independent laboratories across Europe (**Figure 4A** and **Table 1**). Each institution received the same general protocol for the implementation of the amplicon-based genotyping method (18). Nonetheless, the specific setup (e.g. sample collection, RNA extraction, quantitative reverse-transcription PCR (RT-qPCR) test, choice of replicate samples) was left to the discretion of each laboratory. The raw sequencing data for 227 clinical samples (**Table 1**) was processed using the same analysis workflow to ensure comparable results.

**Figure 4.**
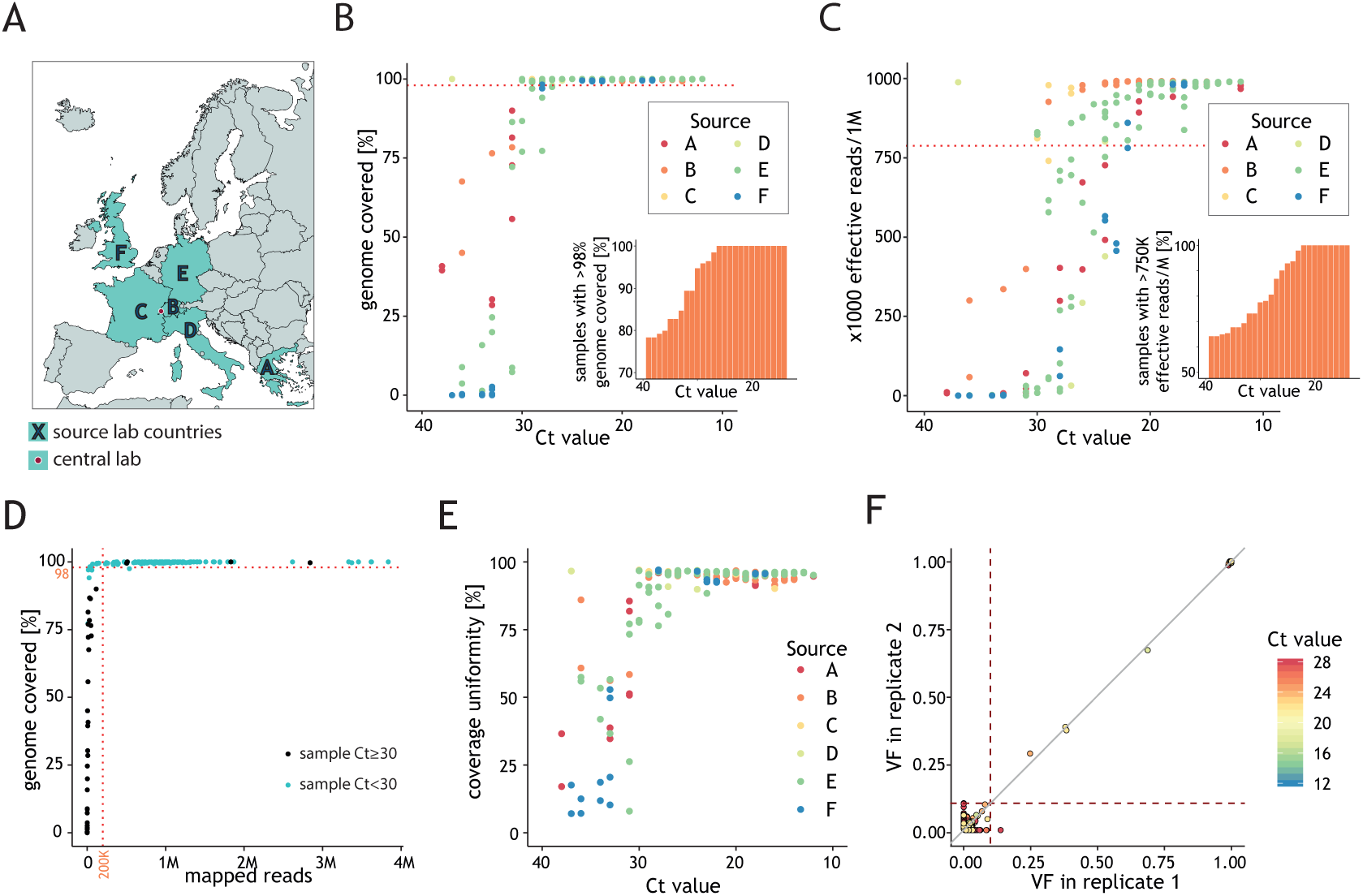
Viral genotype assignment in clinical samples reflects global genome diversity. (A) The multicentre study involved 6 laboratories, located in different European countries, which generated datasets analyzed at a central location (SOPHiA Genetics, Switzerland). (B) Fraction of viral genome covered by at least 10 reads (y-axis) as a function of the Ct value (y-axis). Each point represents the results for a sample, colour coded according to the source lab. The dashed line indicates 98% coverage breadth. The percentage of samples with at least 98% genome coverage breadth (y-axis) below a given Ct (x-axis) is represented in the inset. (C) The number of reads mapping to the SARS-CoV-2 genome per 1M total reads (y-axis) as a function of the Ct value of the clinical samples (x-axis). Each point represents the results for a sample colour coded according to the source lab. The dashed line indicates 750K mapped reads/M. The percentage of samples with at least 750K reads per million mapped reads (y-axis) below a given Ct (x-axis) is represented in the inset. (D) Fraction of viral genome covered by at least 10 reads (y-axis) as a function of the number of reads mapping to the SARS-CoV-2 genome for the clinical samples (x-axis). Each point represents a sample and is coloured blue or black if Ct value is below or higher-or-equal to 30, respectively. The horizontal dotted line indicates 98% coverage breadth and vertical dotted line indicates 200K mapped reads; points are coloured according to the sample Ct value. (E) Percentage of genome coverage uniformity (y-axis) as a function of input of the sample Ct value (x-axis). Each point represents the results for a sample colour coded according to the source lab. (F) Relationship between variant fraction for variant calls in clinical samples processed in replicates (refer to Table 1 for details) and with genome coverage breadth >98%. Dotted lines demarcate VAF=0.1. Variants are coloured based on the Ct value of the replicate.

**Table 1.**
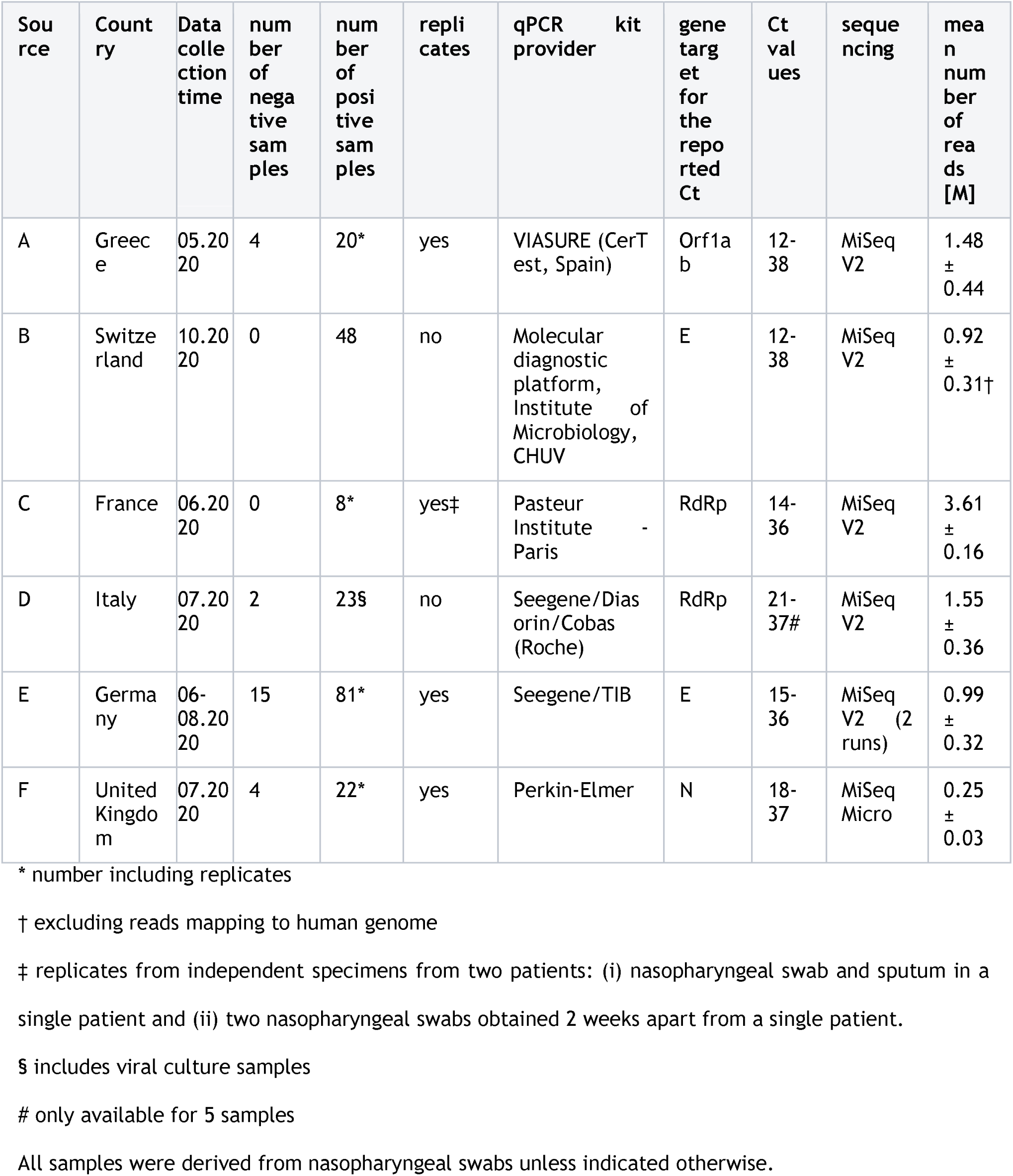
Summary of clinical datasets used in this study.

In the clinical setting, the RT-qPCR cycle threshold (Ct) value, which correlates with the number of viral copies (**Figure S4A, Additional File 1**), is commonly used as a proxy for the number of SARS-CoV-2 genomes in the sample. For the samples included in this study the Ct values varied between 12 and 38 (median Ct = 23, **Figure S4B, Additional File 1**). These values were provided by the source laboratory and obtained using distinct RT-qPCR assays, which we anticipate should maximize the universal applicability of our conclusions (**Table 1**). To investigate the impact of the method and operator on Ct estimates, samples containing 1-100’000 SARS-CoV-2-C2 g.c.p.r. were analysed by RT-qPCR in two independent labs using different assays (30,31). We found that the Ct values obtained from these samples using independent methods are highly correlated (**Figure S4A, Additional File 1**) as previously found (32). Calibration results suggest that samples with 1000 g.c.p.r. should generally yield Ct values in the range of 29-31, consistent with previous reports (30-32). Accordingly, 96% of clinical samples with Ct of less than 29 had at least 98% genome coverage breadth (99.6% on average) (**Figure 4B** and **S4C, Additional File 1**) and 81% yielded at least 750K mapped reads per million (**Figure 4C**).

Similarly to what was observed in the experiments with synthetic RNA, no improvement in the genome coverage breadth was observed above the recommended value of 200K mapped reads for these samples (**Figure 4D**). For samples with Ct lower than 26, we typically obtained between 70-90% of mapped reads. For these samples we recommend sequencing to a depth of ~280K reads to ensure sufficient genome coverage breadth and depth. Samples with Ct values between 26 and 30 display generally good coverage breadth (**Figure 4B**) and uniformity (**Figure 4E**) but variable fraction of effective reads (**Figure 4C**) and therefore should be sequenced at a higher depth to achieve the recommended 200K mapped reads and 98% coverage breadth cut-offs.

Based on our analysis of synthetic RNA, we recommend that samples with at least 200K mapped reads and >98% coverage breadth should be used for the detection of variants (VAF ≥ 0.1) in the SARS-CoV-2 genome. To confirm the validity of this limit for clinical samples, we took advantage of replicates present in our dataset and compared variant allele fractions measured across the replicates with more than 200K mapped reads and at least 98% coverage breadth (n=13 sample pairs) (**Figure 4F**). Most variants (99.2%, 126/127) with VAF >0.1 were reproducibly detected and their VAF values were strongly correlated (Pearson r=0.996). In contrast, for variants detected in at least 1 replicate at a VAF below 0.1 no correlation between replicates was detected (r=-0.45). The elevated number of low frequency variants in samples that do not fulfill our recommendations suggests the presence of false-positive calls due to increased background noise resulting from lower quantity/quality of the viral RNA (**Figure S4D** and **S4E, Additional File 1**). We conclude that for samples with sufficient viral load the variant calling is reliable for alleles present at a fraction of at least 0.1.

### Viral genotype assignment in clinical samples reflects global genome diversity

One of the main goals of SARS-CoV-2 genotyping is to determine the viral strain present in the sample. We performed variant calling in all unique clinical samples. We assumed that variants with VAF of 0.9 or higher were present in all viral genotypes of the sample and referred to these as ‘clonal’ (**Figure 5A**). The remaining alleles (0.1≤VAF<0.9) were labelled as ‘minor’ and are likely the result of co-infection by more than one virus genotype or intra-host mutations (‘quasi-species’).

We investigated whether minor variants were more likely to result from infection by multiple viral strains or else were the consequence of intra-host mutations. We estimated the pairwise sequence distance between contemporaneous samples isolated in the same lab to account for the impact of time-related divergence increase and differences in viral genotypes present in different European countries (33) (**Figure S5A, Additional File 1**). The distribution of the expected number of inter-sample clonal variant differences was compared to the distribution of minor alleles observed in each sample, which is an estimate of intra-host genome diversity (**Figure S5B, Additional File 1**). The number of differences observed between isolates (mean=10, mode=2) was higher than that observed at intra-sample level (mean=2.8, mode=0). This observation supports that intra-host diversity is more likely to arise from viral mutations following host infection (quasi-species) than from infection by multiple strains.

**Figure 5.**
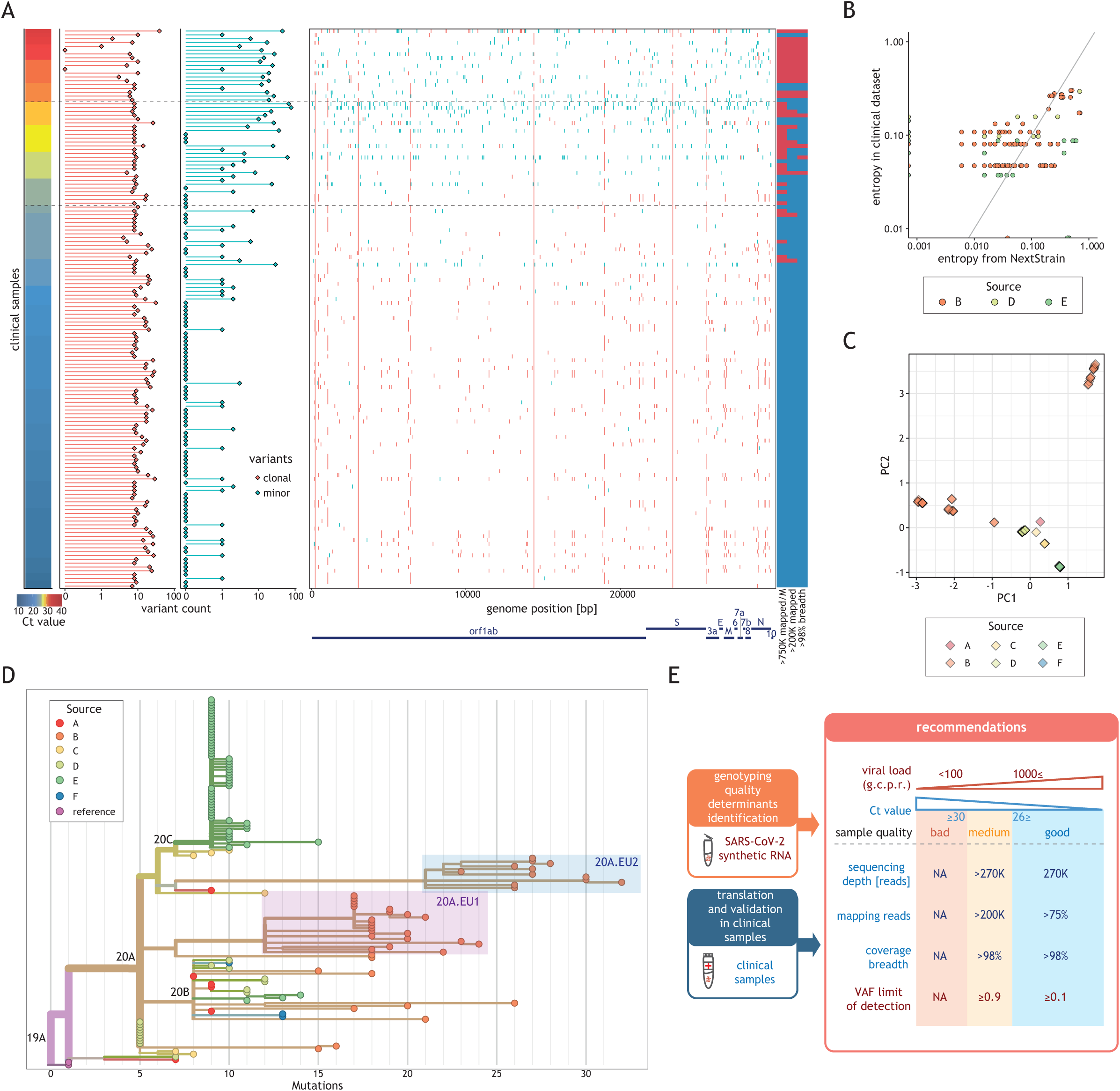
Variant frequencies found in the clinical dataset reflect global frequencies. (A) Summary of the variant calling analysis for all unique clinical samples (rows) sorted by the Ct value (left). The horizontal lines indicate Ct values of 26 and 30. The number of clonal (VAF≥0.9, red) and minor (0.1<VAF<0.9, cyan) variants for each sample is represented as horizontal bar-plots (middle left). The position of each clonal (red) and minor (cyan) variant is displayed along the genome (middle right). Classification of the samples relative to the different recommendation (listed below the column); blue indicates the recommendation was fulfilled and red that it was not (right). (B) Relationship between the entropy estimated for all clonal variants in clinical samples (y-axis) and the entropy of the same variants in samples collected in the same country and during the same period according to Nextstrain (33) (x-axis). Only samples with >200 K effective reads and 98% coverage breadth from centres with data for more than 15 samples were considered in this analysis. (C) 2-D principal component analysis results of clonal variants in clinical isolates (points). Points are coloured based on the sample source. (D) Phylogenetic tree of all clinical isolates with >200 K effective reads and 98% coverage breadth criteria. Samples are coloured according to the source. Clades (according to Nextstrain) are indicated. Samples corresponding to subclade 20A.EU.1 and 20A.EU.2 are highlighted by red and blue boxes, respectively. Length of the branches reflects the number of mutations (x-axis). The tree visualization was generated using the Nextstrain platform. (E) Schematic representation of the recommendations for reliable genotyping with amplicon-based approach. We used synthetic viral genomes to determine the minimal viral load and VAF. We validated these recommendations and made them broadly applicable using clinical samples by determining the minimal sequencing depth, fraction of mapped reads and coverage breadth. Samples were classified into three quality categories based on their viral load: good (≥1000 g.c.p.r.), intermediate (uncertain g.c.p.r., Ct values in the range 26-30) and bad (<100 g.c.p.r., typically value Ct>30).

The majority (133/135, 98.5%) of clinical samples we obtained from the various European centers was characterized by the same 4 alternative alleles (at positions 241, 3037, 14408, 23403) (**Figure 5A**). These variants, along with the fifth most frequent variant (position 25563), are repeatedly reported in global datasets (11). The frequencies of clonal variants found across these clinical datasets are similar to frequencies expected from samples collected during the same period at the respective location, as illustrated by correlation with Shannon entropy values provided by NextStrain (33) (Pearson r=0.701 and p<10^-15^) (**Figure 5B**). In addition, principal component analysis supports that some variants are common within samples from specific sources, likely reflecting the relationship between genetic diversity and the location and the time of collection (**Figure 5C**). A large fraction of the variability is explained by the difference between samples collected before and after July (**Figure S5C, Additional File 1**). Almost all clinical samples collected before July 2020 belong to clade 20A, characterized by the presence of variant D614G that seeded the SARS-CoV-2 outbreak in Europe, and its daughter clades 20B and 20C (**Figure 5D**). The increased diversity in clonal variants observed after July 2020 is explained by the presence of 81.8% (36/44) of clinical isolates belonging to subclades 20A.EU1 and 20A.EU2. These two strains are thought to have emerged in Europe during early summer 2020 and their spread across multiple European countries facilitated by increased cross-border travelling during the summer holiday (34). The above results support the validity of clinical sample genotypes determined using our recommendations.

## Discussion

Monitoring of global and local disease spread during the SARS-CoV-2 pandemic requires fast, sensitive and precise methods for pathogen genotyping. Amplicon-based approaches are currently the most widely used method in this context (16-21), yet their limits of performance have not been systematically evaluated. Consequently, the quality of the genomes obtained cannot be easily assessed, resulting in errors that are difficult to trace and submission of viral genome sequences of poor quality to public repositories (11-14). Here, we used both synthetic RNA and clinical isolates processed by multiple clinical laboratories to dissect the factors which need to be taken into account to obtain reliable SARS-CoV-2 genome information using an amplicon-based approach (**Figure 5E**).

Variant calling accuracy strongly depends on the quality and quantity of starting material because the noise level - understood as incorrect base-calls due to a combination of RT, PCR and sequencing errors - is inversely correlated with the number of viral genomes and the sequencing depth. Our analysis of synthetic genome controls shows that variants at VAF>0.1 can be reliably detected in samples with at least 1000 g.c.p.r., which corresponds to approximately 100 viral genomes per ml. Libraries generated from these samples yield at least 750K mapped reads per million sequenced reads and are sufficiently complex to ensure at least 98% of genome coverage at a depth of 10 reads per nucleotide when at least 200K reads are mapped.

We used viral sample sequencing data from a multi-centre collaboration across Europe to investigate how these recommendations can be implemented and what their impact is on clinical sample genotypes. We show that the fraction of effectively mapping reads and the breadth of coverage can be used to evaluate the quality and quantity of the original input material in diverse clinical samples. Based on the qPCR calibration and the genome coverage breadth comparison we estimate that the viral load of >1000 g.c.p.r. generally corresponds to samples with qPCR Ct value lower than 30. In clinical samples with Ct values between 26 and 30, the fraction of effective reads is highly variable likely due to various technical factors. An increase in the sequencing depth might improve coverage breadth, depth and uniformity for these samples. This does not apply in cases where the genome is highly degraded or low abundance (i.e. Ct>30). Our results are in agreement with recent reports showing poor genome coverage for the majority of samples with Ct>30 (28,35,36). Importantly, our conclusions hold true irrespective of the source lab and the specific RT-qPCR conditions, ensuring the general applicability of our recommendations.

Despite being frequently used in clinical epidemiology (34), the variable quality of sequences deposited in public repositories is hard to infer and will impact downstream analyses. Using our recommendations and considering only variants at frequencies higher than the estimated limit of detection of 0.1, we determined genotypes of the clinical samples from several sources. We found that as expected, virtually all clinical samples have the D614G variant characteristic of the SARS-CoV-2 strain at the origin of the European outbreak. We also detected source-specific polymorphisms and the presence of representative samples from the main European clades. These include 2 strains that are thought to have emerged early in the summer of 2020 and spread across Europe by tourists travelling to, and from, their holiday destination, specifically in samples collected in October 2020.

Even if several studies report variants found at frequencies below 0.1 (25,27,37-39) only a few evaluated the confidence of such calls (26,40). We found that the number of variants with VAF>0.1 detected in samples with at least 1000 g.c.p.r is similar between contemporaneous samples and increases with time since the start of the outbreak, as expected. In contrast, in clinical samples with low viral load we observed an elevated number of low frequency variants, in line with increased false positive calls due to the presence of elevated noise signal. These observations illustrate the risk of not considering the impact of technical factors on the accuracy of the calls in SARS-CoV-2 genotyping and are a testament to the value of applying clear guidelines to select samples of sufficient quality to inform genomic epidemiology studies.

## Conclusions

After careful evaluation of multiple factors affecting SARS-CoV-2 amplicon-based genotyping, we demonstrate that at least 1000 viral copies per reaction should be used for reliable detection of variants with VAF > 0.1 using amplicon-based approaches. We show that variants found in clinical isolates with viral loads above this threshold allow to distinguish samples based on their geographical origin and time of sampling, as expected. Widespread implementation of technical guidelines for SARS-CoV-2 genotyping will improve the quality of reported genotypes and the reliability of downstream analysis.

## Methods

### Library preparation with synthetic viral genome and sequencing

Three versions of Synthetic SARS-CoV-2 genomes – Control 2 identical to the reference genome (NCBI NC_0455l2.2/Genbank ID MN908947.3, #102024), Control 1 containing four variants (3 SNVs and a 10 bp-long deletion, MT007544.1, #102019)) and Control 4 containing 7 SNVs (MT106054.1, #102862:) - purchased from Twist Bioscience were used as input, at indicated number of copies. Individual sample information is summarized in **Table S1, Additional File 2**. The number of viral genome copies for the strain specified in the experiment was spiked into 50 ng of Universal Human Reference RNA (ThermoFisher Scientific #QS0639) to the final volume of 11 μl. Libraries were prepared with CleanPlex SARS-CoV-2 Panel (Paragon Genomics #918011) according to manufacturer’s instructions (18). The resulting libraries were quantified using Qubit 1X dsDNA HS Assay (ThermoFisher Scientific, #Q33231) and library size was estimated by capillary electrophoresis. Libraries were mixed at equimolar amounts, with 5% PhiX and 150bp-long paired-end reads sequenced using Illumina MiSeq or NextSeq sequencers.

### Clinical samples

Clinical data was obtained from six independent European institutions (**Table 1**). Only samples where the presence of SARS-CoV-2 could be confirmed by qPCR were considered. For a subset of clinical samples samples we had either technical (n=35) or biological (n=2) replicates. Sample information is summarized in **Table S1, Additional File 2.** All samples were processed with CleanPlex SARS-CoV-2 Panel (Paragon Genomics #918011) using either 25 or 50 ng total RNA as input. Resulting libraries were quantified, evaluated by capillary electrophoresis and 150 bp-long pair read sequenced using Illumina sequencers by the source lab. Ct values were provided by the source lab. For each sample we approximated the reported Ct to the closest integer to facilitate comparisons.

### Data processing

Sequencing reads alignment to the reference genome NC_045512.2 and variant calls were performed with SOPHiA Genetics proprietary pipeline. Mapping to human transcriptome and genome were performed using STAR (ver 2.7) (41) and bwa mem (version 0.7.12-r1039) (42), respectively. Read coverage was estimated using bedtools coverage from the bedtools suite (version v2.29.2) (43) for the portion of the viral genome targeted by the amplicons (positions 36-29,844 of the reference genome). Genome coverage depth and breadth were calculated for each sample. Coverage uniformity is defined as the fraction of genome covered at a depth in the range between the 20% and 500% of the median depth. Read subsampling was performed with samtools v1.9 (44).

### Determination of VAF limit of detection for intra-host variability

Synthetic viral genome controls (SARS-CoV-2-C2, SARS-CoV-2-C1 and SARS-CoV-2-C4) were mixed to achieve the desired fraction (0-0.2) and spiked into 50 ng of human reference RNA at different defined viral load between 100-10’000. Libraries were prepared, sequenced and analysed as previously described. Variants called in the replicates were pooled for downstream analysis. For a given VAF threshold all known variants detected above or below the expected VAF were considered to be True Positives or False Negatives, respectively. Background variants were considered to be either False Positives or True Negatives, when their VAF was above or below the tested threshold, respectively. We used true/false positive/negative calls at different VAF to estimate the area under Receiver Operating Characteristic (ROC) curve (AUC), sensitivity and specificity with different number of mapped reads and at different VAF.

### Analysis of intra-host and inter-host diversity

In case of replicates only one of the samples, chosen at random, was included. Variants called with a frequency between 0.01 and 0.9 were labelled as minor and variants called with a frequency of at least 0.9 were labelled as clonal. Only clinical samples that met the criterion of >98% genome coverage breadth and at least 200K mapped reads were retained for the downstream analyses (n=135). We considered samples from each individual source separately and calculated the number of all possible clonal pairwise differences for all possible sample pairs. We compared the distribution of all possible pairwise differences to per-sample counts of minor variants.

In order to compare diversity within our clinical dataset to that reported in NextStrain (33), we calculated per-position Shannon entropy individually for datasets with at least 15 samples. These values were then compared with values reported in NextStrain for a country and sampling period matching the respective dataset. As the precise sampling date was not available for most of the samples, we chose the time period since December 2019 until the month preceding the month of data upload.

### Phylogenetic anaylsis

Phylogenetic analysis of clonal genomes from clinical samples was performed using the Nextstrain pipeline with default settings (https://github.com/nextstrain/ncov/) (33).

### RT-qPCR calibration

RT-qPCR calibration was performed in two institutions. Source B performed the test using two types of reference material: synthetic SARS-CoV-2 RNA or plasmid bearing viral genes using the test described in (45). SOPHiA Genetics lab performed the test using synthetic SARS-CoV-2 RNA, CDC-USA assay targeting gene N (IDT # 10006713) and One Step PrimeScript™ III RT-PCR Kit (TaKaRa #RR600A). Serial dilutions of reference material were prepared ranging from 1 to ~10M genome equivalents per reaction. RT-qPCR was performed according to manufacturer’s instructions.

### Statistical analysis

Statistical data analyses were performed using the R software environment for statistical computing (46) and graphics with the ggplot2 package (47).

## Declarations

### Ethics approval and consent to participate

Not applicable

## Consent for publication

Not applicable

## Availability of data and materials

The datasets generated with synthetic SARS-CoV-2 genome and analysed during the current study are available in the Sequence Read Archive (SRA) repository under accession number SUB8654793. The data generated with clinical samples and used in this study are available from each respective source institution (listed in Table 1) but restrictions apply to the availability of these data, which were used under license for the current study, and so are not publicly available. Requests must be directed to the owner of each dataset.

## Authors’ contributions

**Study conception and design:** S.K, A.C.M., A.W, and Z.X; **Acquisition of data:** S.K, J. S., C. B., F. D.M., S. P., T. B., Y. D., C.A., H.S., A.S., M. M., E.W.S., A.S.M., J. C., R. S., P. C., M. S., G. G. and C. T.; **Analysis and data interpretation:** S.K., X.X., Y.W. and L.S; **Study Supervision:** M. P., A.W. and Z.X.; **Drafting of manuscript:** S.K., A.C.M., L.M.S., V.P., and Z.X. All authors read and approved the final manuscript.

## Acknowledgements

We would like to thank Dr. Frederic Michaud for discussion on how to visualise viral diversity; Dennis Deschka (LABCON-OWL GmbH, Bad Salzuflen) for his excellent advise and technical support during the project; Sébastien Aeby and Dr. Katia Jaton (Institut de Microbiologie, CHUV) for their help in the acquisition of clinical sample data; Pr. Christelle Thauvin and Pr. Laurence Faivre (département de génétique, CHU Dijon); Hélène Giraudon, Catherine Manoha (Laboratoire de Virologie, CHU Dijon); Pr Lionel PIROTH (Département des Maladies Infectieuses et Tropicales CHU Dijon); Pr Etienne Carbonnelle & Dr Ségolène Brichler (Service de Microbiologie, CHU Avicenne) for their help in aquisition of clinical sample data; Dr. Stavroula Samara and Dr. Katerina Oikonomaki for help in library preparation.

## Authors’ information

L.M.S. contributed to this publication as a co-founder of SOPHiA GENETICS and not part of his Stanford University duties or responsibilities.

---

**Figure S1.**
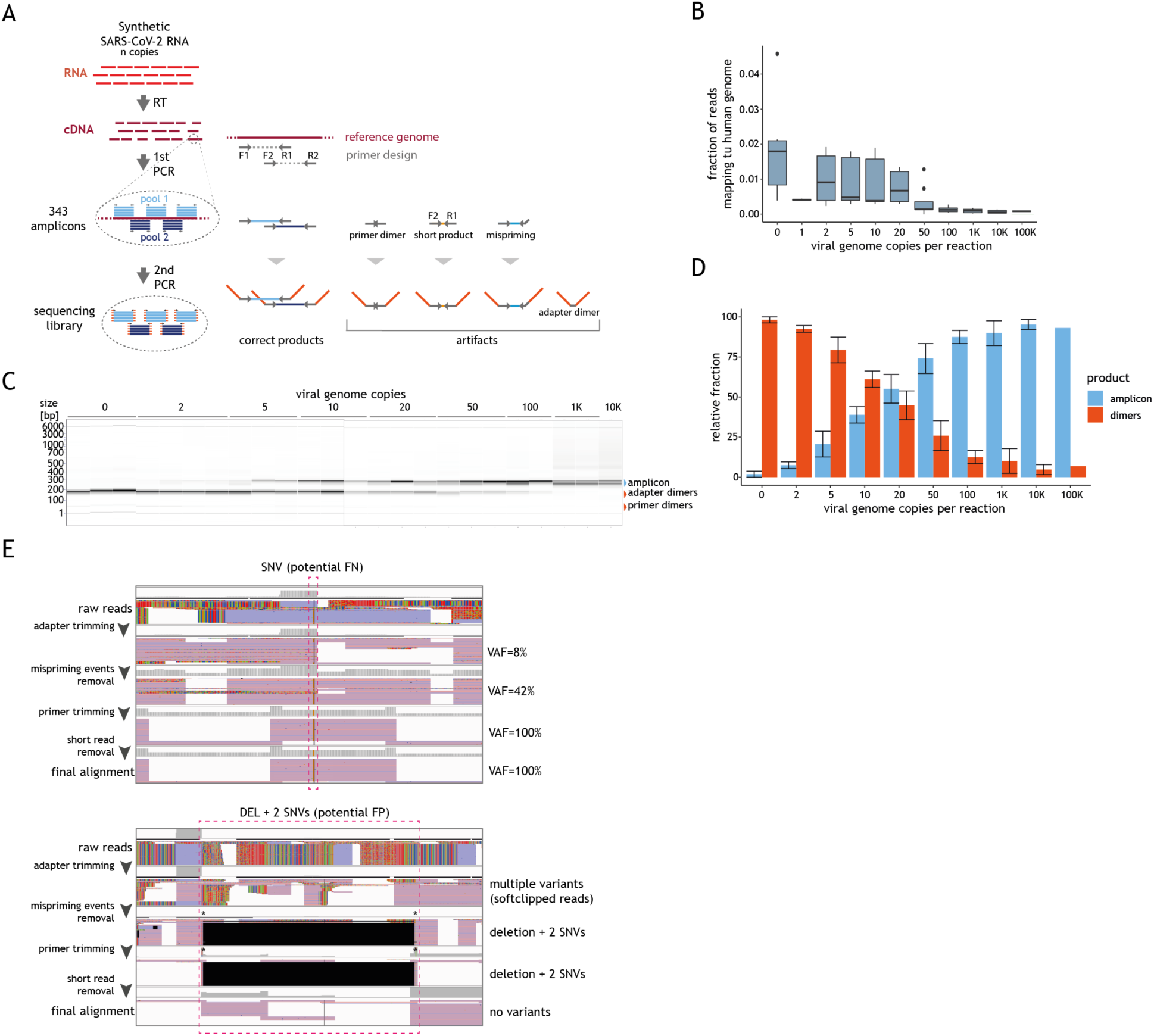
(**A**) Schematic representation of amplicon-based workflow and the source and type of artifacts it can introduce in the sequencing data. The RNA is converted into cDNA and amplified, in two pools, into 343 partially overlapping amplicons (example primer design F1-R1 and F2-R2 shown). After pooling, the amplicons are subsequently converted into sequencing libraries. These two PCR steps can introduce different types of artifacts as illustrated by the panel on the right. (B) Distribution of the fraction of sequencing reads aligning to human genome (y-axis) for samples with varying amounts of synthetic viral genome (x-axis). The horizontal line in the boxplot indicates the median and the whiskers the 5% and 95% quantile. (C) Capillary electrophoresis profiles of libraries obtained for samples with varying amounts of synthetic SARS-CoV-2 genomes (indicated on the top of the lane). Fragment size based on control DNA ladder is indicated on the left and the expected size of SARS-CoV-2 amplicons (red arrow) or artifacts (blue arrow) is indicated on the right. (D) Fraction of DNA fragments with sizes corresponding to SARS-CoV-2 amplicon or artifact quantification from the electrophoretic library profiles obtained for samples with varying amounts of synthetic SARS-CoV-2 genomes. Bars represent average of at least 3 replicates for each amount and whiskers the standard deviation. (E) Genome browser view examples of artifact containing reads and their impact on inferred genotype before and after their removal; top panel depicts a SNP initially detected only at a low VF (8%, below the cut-off for being reported in the final genome sequence); bottom panel shows reads supporting 2 SNPs and a deletion which are not present in the final alignment.

**Figure S2.**
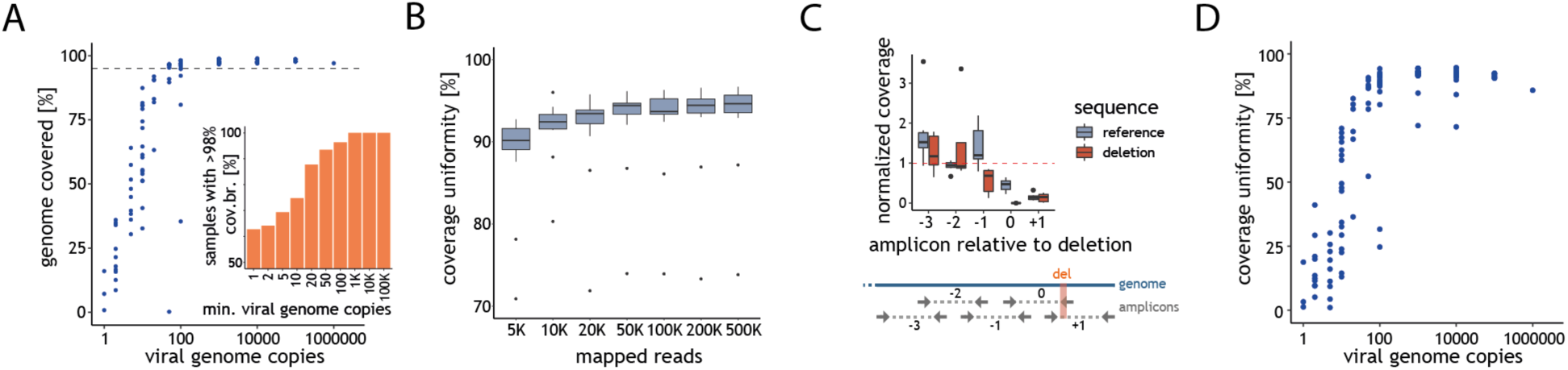
(A) Coverage breadth (y-axis) as a function of the number of synthetic viral genomes in the input material (x-axis). Each point represents the data for one sample. Inset depicts the fraction of samples with coverage breadth >98% as a function of the synthetic viral genomes in the input material (x-axis, as g.c.p.r.) (B) Distribution of the coverage uniformity (y-axis) as a function of the number of mapped reads (x-axis). The horizontal line in the boxplot indicates the median and the whiskers the 5% and 95% quantile. (C) Distribution of the median-normalized coverage at all amplicons spanning (0, +1) or neighbouring (−3,−2 and −1) the deletion present in SARS-CoV-2-C1 in samples containing copies of this viral control (deletion, red) and or SARS-CoV-2-C2 (reference, blue). The horizontal line in the boxplot indicates the median and the whiskers the 5% and 95% quantile. The schematic representation of the different amplicons relative to the deletion SARS-CoV-2-C1 is depicted on the lower panel. (D) Percentage of the coverage uniformity (y-axis) as a function of the number of synthetic viral genome copies in the sample (X-axis, g.c.p.r.). Each point represents the data for one sample.

**Figure S3.**
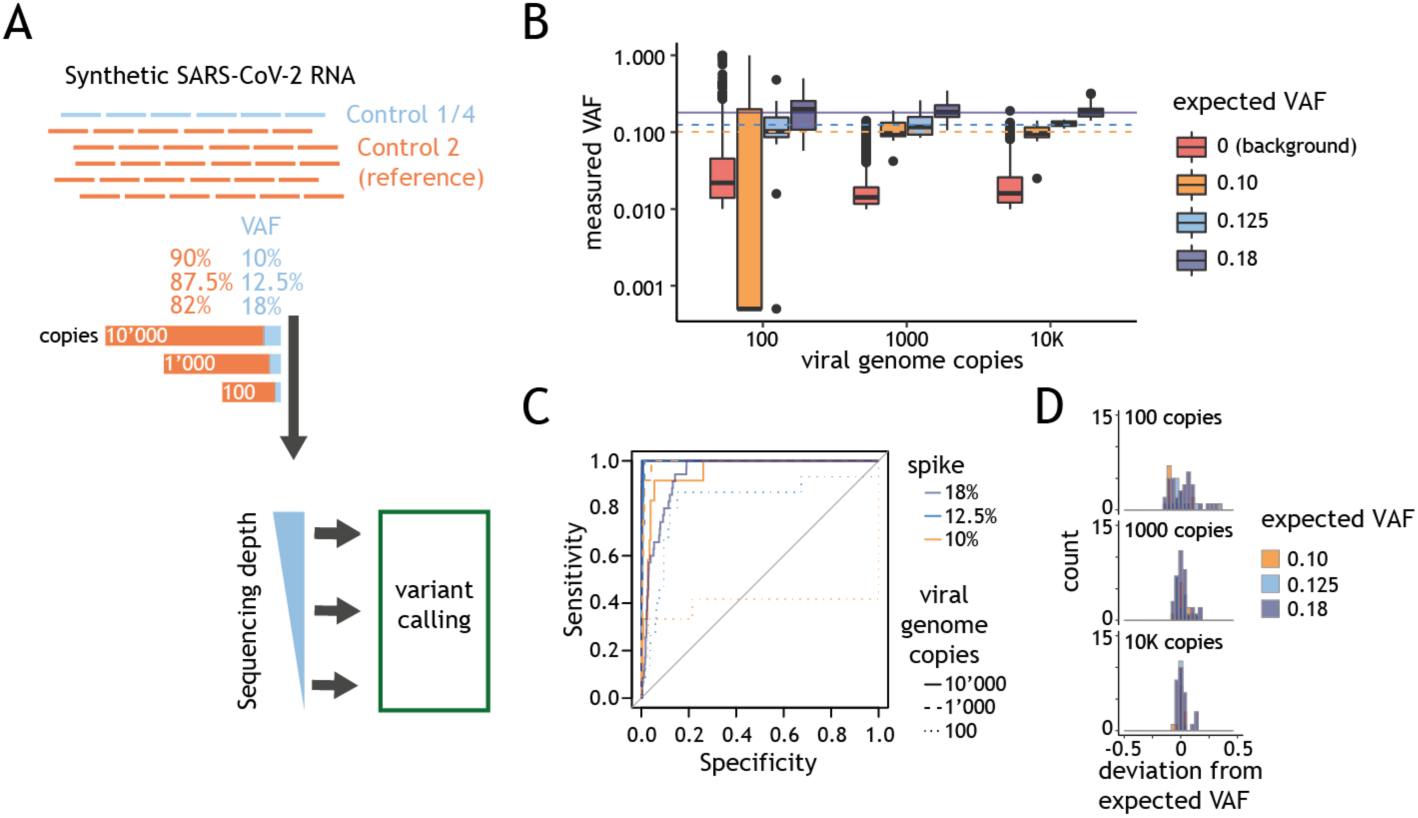
(A) Schematic representation of the experimental design for the determination of the impact of intra-host variability on variant calling performance. Varying amounts of SARS-CoV-2 Control 1 or 4 (blue) were mixed with SARS-CoV-2 synthetic genome reference (Control 2, orange) to obtain desired VAFs and varying viral genome copy mixes (between 100, 1000 and 10000 g.c.p.r.) spiked into human RNA. Variant calling was performed for samples sequenced at varying depth. (B) Distribution of the variant fraction for known variants (y-axis), expected fraction 0.1, 0.125 and 0.18 (yellow, blue and purple) or background call (red), as a function of the viral genome copies in the input material. The horizontal line in the boxplot indicates the median and the whiskers the 5% and 95% quantile (C) Sensitivity (y-axis) as a function of the specificity (x-axis) for variants at different VAF. The ROC curves for are colour coded depending on expected VAF of the variants analysed. Data for samples with 100, 1000 and 10 000 viral copies are represented as dotted, dashed or straight lines, respectively. (D) Histogram representing the number of variants (y-axis) displaying a given deviation between measured and expected VAF (x-axis) for samples with varying viral genome amounts (g.c.p.r).

**Figure S4.**
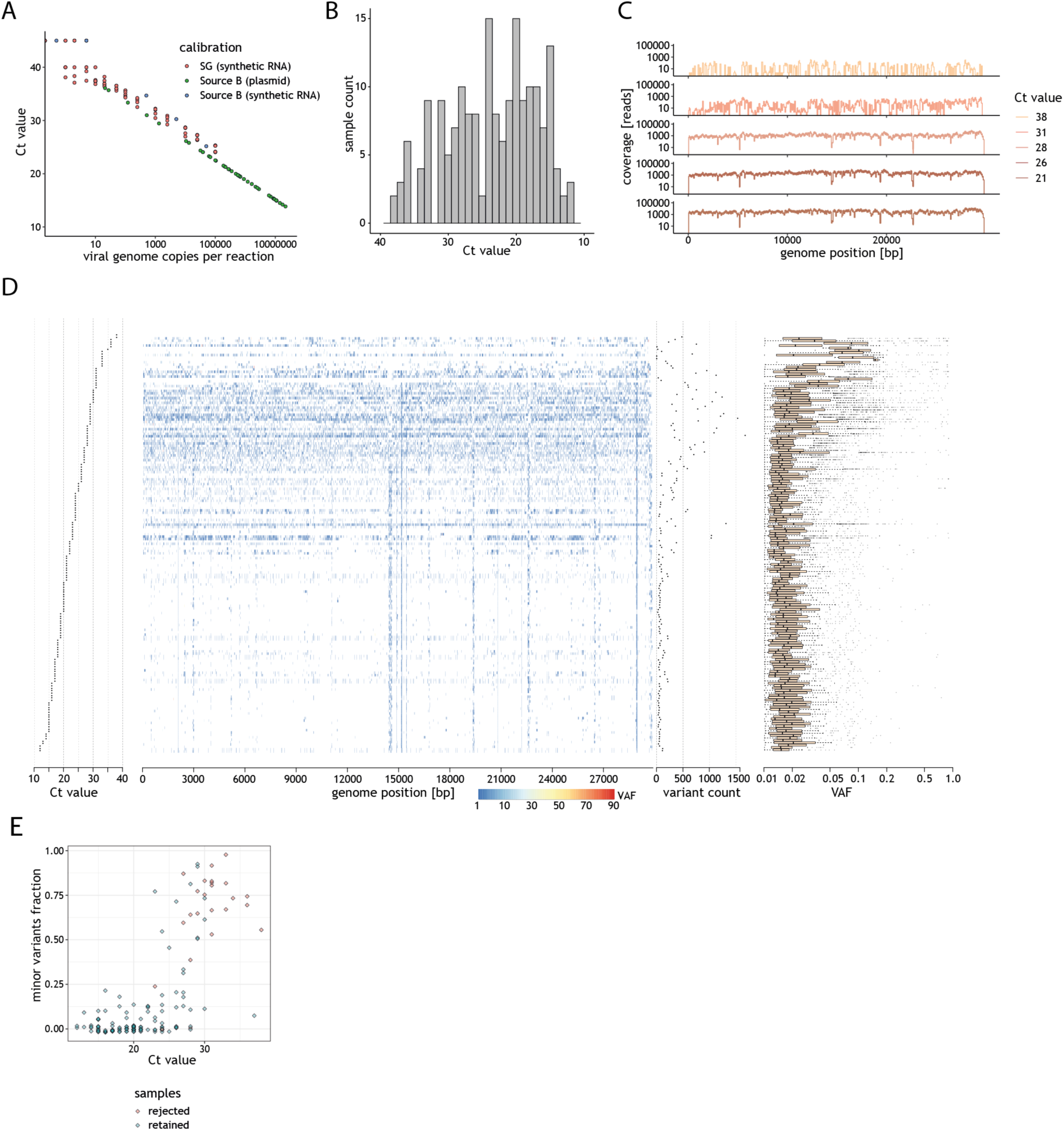
(A) Relationship between the Ct value (y-axis) and the viral genome copies based either on synthetic SARS-CoV-2 RNA or plasmids encoding viral genes, performed by two independent laboratories (SOPHiA Genetics (SG) or Source B). (B) Distribution of Ct values for all clinical samples where the value was provided. (C) Ideograms depicting the genome coverage (Y-axis) for representative samples with varying Ct values. (D) Summary of the variant calling analysis, taking into account all VAF values, for all unique clinical samples (rows) sorted by the Ct value (left); the position of each variant (blue) is displayed along the genome (central panel). Right panels represent the variant count and distribution of the VAF of all variants detected in a given sample. (E) Fraction of variants with 0.1<VAF<0.9 (minor variants, y-axis) as a function of the sample Ct (x-axis). Each point corresponds to the results of one clinical sample. Samples that fulfil the recommendations (>200K effective reads and 98% coverage breadth) are coloured in blue. The remaining samples are represented by points coloured red.

**Figure S5.**
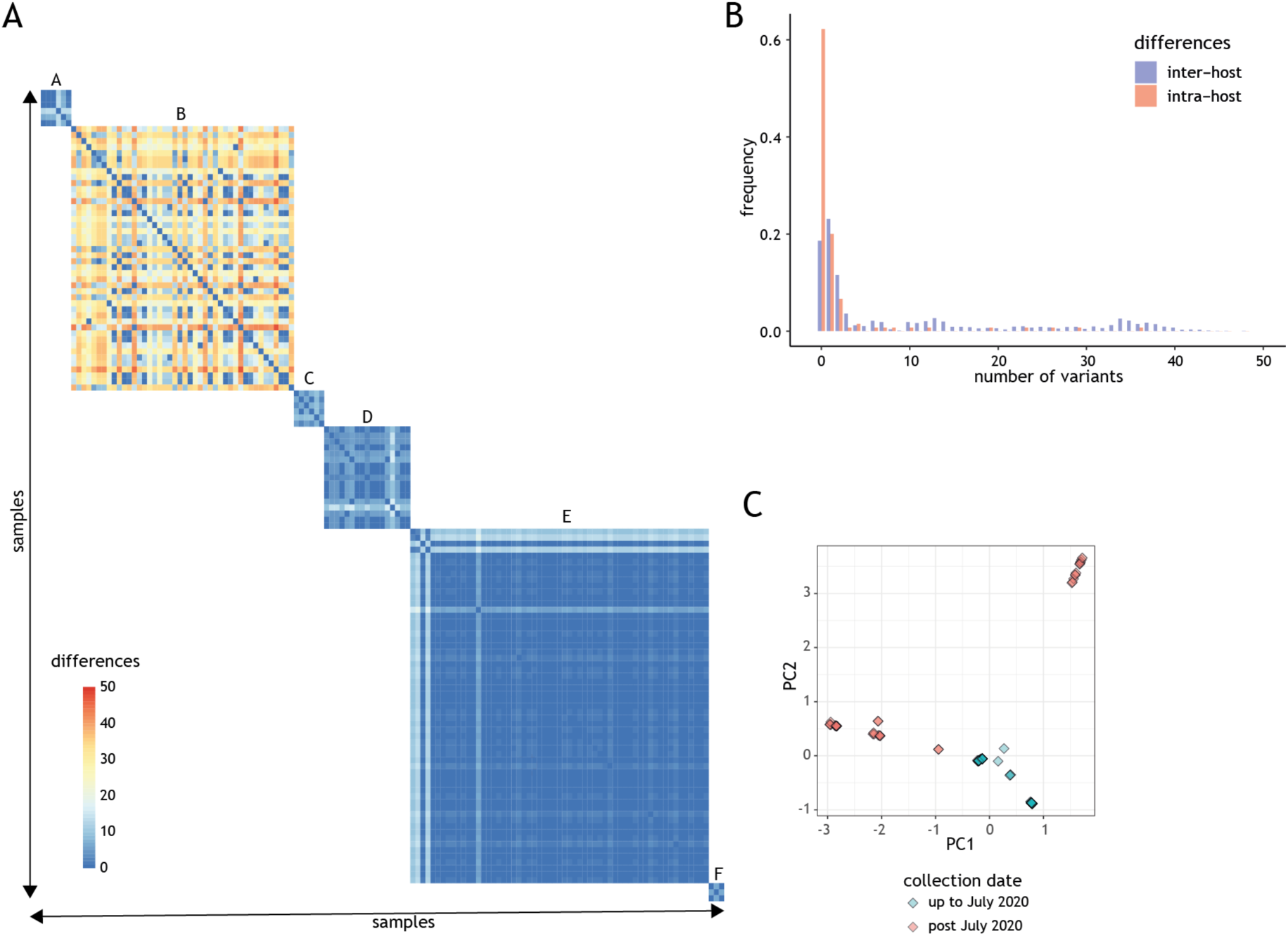
(A) Heatmap of the number of clonal differences between contemporaneous clinical isolates collected and processed by the same centres (sources A-F). (B) Histogram representing the distribution of the inter-(red) and intra-sample (blue) diversity (as number of variants). (C) 2-D principal component analysis results of clonal variants in clinical isolates (points). Points are coloured based on the data collection date (up to July 2020 or after).

